# Telomerase reverse transcriptase is required for resistance to mycobacterial infection

**DOI:** 10.1101/2025.05.15.654386

**Authors:** Darryl JY Han, David M Costa, Geyang Luo, Kathryn Wright, Manitosh Pandey, Denise Wee, Yao Chen, Semih Can Akincilar, Vinay Tergaonkar, Rajkumar Dorajoo, Guillaume Carissimo, Amit Singhal, Philip M Elks, Pamela S Ellis, Catarina M Henriques, Bo Yan, Serge Mostowy, Stefan H Oehlers

## Abstract

Age is an important risk factor for infections such as tuberculosis (TB). Telomerase is expressed in immune cells yet leukocyte telomere length declines during ageing suggesting an age-dependent loss of telomerase activity in the immune system. Leukocyte telomere length has been correlated with worse outcomes in TB patients, however the mechanisms linking telomere biology to TB susceptibility and response to therapy are unexplored. Here we use the zebrafish-*Mycobacterium marinum* model to investigate the role of telomerase in TB resistance. We find depletion and inhibition of Tert, the catalytic subunit of telomerase, increases bacterial burden in zebrafish embryos. The Tert depletion infection susceptibility phenotype could not be rescued by p53 or STING depletion suggesting a non-canonical role for Tert in controlling mycobacterial infection. Consistent with a previously described role for Tert in developmental hematopoiesis, we find Tert is necessary for demand-driven emergency myelopoiesis to support containment of extended mycobacterial infection. Our findings establish a previously undescribed role for host telomerase in supporting infection demand-driven hematopoiesis to control infection.

## Introduction

Ageing is one of the most important risk factors for tuberculosis (TB) susceptibility and severity (*1*). Animal models and clinical measurements have identified inflammageing, a persistent low grade interferonopathy triggered by accumulated DNA damage, and immune anergy, caused qualitatively by declining lymphocyte function and quantitatively by the biased production of myeloid cells, as potential mechanisms underlying age-dependent TB susceptibility (*2, 3*). Recent work across guinea pig and human systems have demonstrated that TB accelerates ageing suggesting that *Mycobacterium tuberculosis* may actively benefit from accelerated host ageing and that interception of this process could yield novel targeted adjunctive therapies for an important at-risk group (*4*).

Leukocyte telomere length (LTL) decreases during ageing and influences immune function, specifically affecting susceptibility to infectious diseases driven by host immunopathology including TB (*5–12*). Some aspects of the relationship between LTL and TB biology may be ethnicity specific as while South African patients with longer LTL are more likely to have a successful response to TB treatment, South East Asian and East Asian patients with longer LTL experienced more treatment side effects (*13–15*). As telomerase activity is required for maintenance of LTL, we hypothesized decreased leukocyte expression of the telomerase reverse transcriptase (Tert) may drive the increased susceptibility of the elderly and those with short LTL to mycobacterial infection.

Expression of *tert* by hematopoietic stem cells is necessary for telomere elongation and likely protects chromosomes during hematopoiesis. There is an increasing body of evidence that Tert and the RNA template *TERC* execute important non-canonical functions in immunity as an inflammation-responsive gene aiding the expression of key hematopoietic differentiation genes (*16–18*). Telomerase-expressing macrophages (T-MACS) have been previously described and loss of telomerase expression has been associated with reduced immune cell-autonomous function (*19, 20*).

Here we utilize the zebrafish-*M. marinum* model of human TB to study the role of Tert in mycobacterial infection. We find that Tert is expressed at the site of infection and loss of Tert is causes reduced control of infection. Tert mRNA supplementation restores infection-induced, demand-driven hematopoiesis, highlighting the role of Tert in supporting hematopoietic resilience during chronic infection.

## Results

### Telomerase reverse transcriptase is expressed at the host-mycobacterial interface

To determine if T-MACs are present in mycobacterial granulomas we analyzed a previously published scRNAseq dataset of dissected zebrafish-*M. marinum* granulomas (*21*). We found *tert* is most expressed in proliferative cells, as expected from canonical function in telomere elongation of proliferating cells and matching the expression of the DNA polymerase subunit *pola2* (Figure 1A). Expression of *tert* was detected in myeloid-derived macrophage subsets, many of which lacked *pola2* expression suggesting a non-proliferative role of *tert* expression. There appeared to be comparable levels of *tert* expression across epithelialized macrophages (Epmacs, Cluster 3, and cdh1+ve) which make up the ring-like structure of mature granulomas compared to the “non-structural” myeloid cells (monocytes, inflammatory macrophages, DCs). The overall level of *tert* expression in the myeloid cell clusters was less than half of the level of *tert* expression proliferating cell clusters.

**Figure 1:**
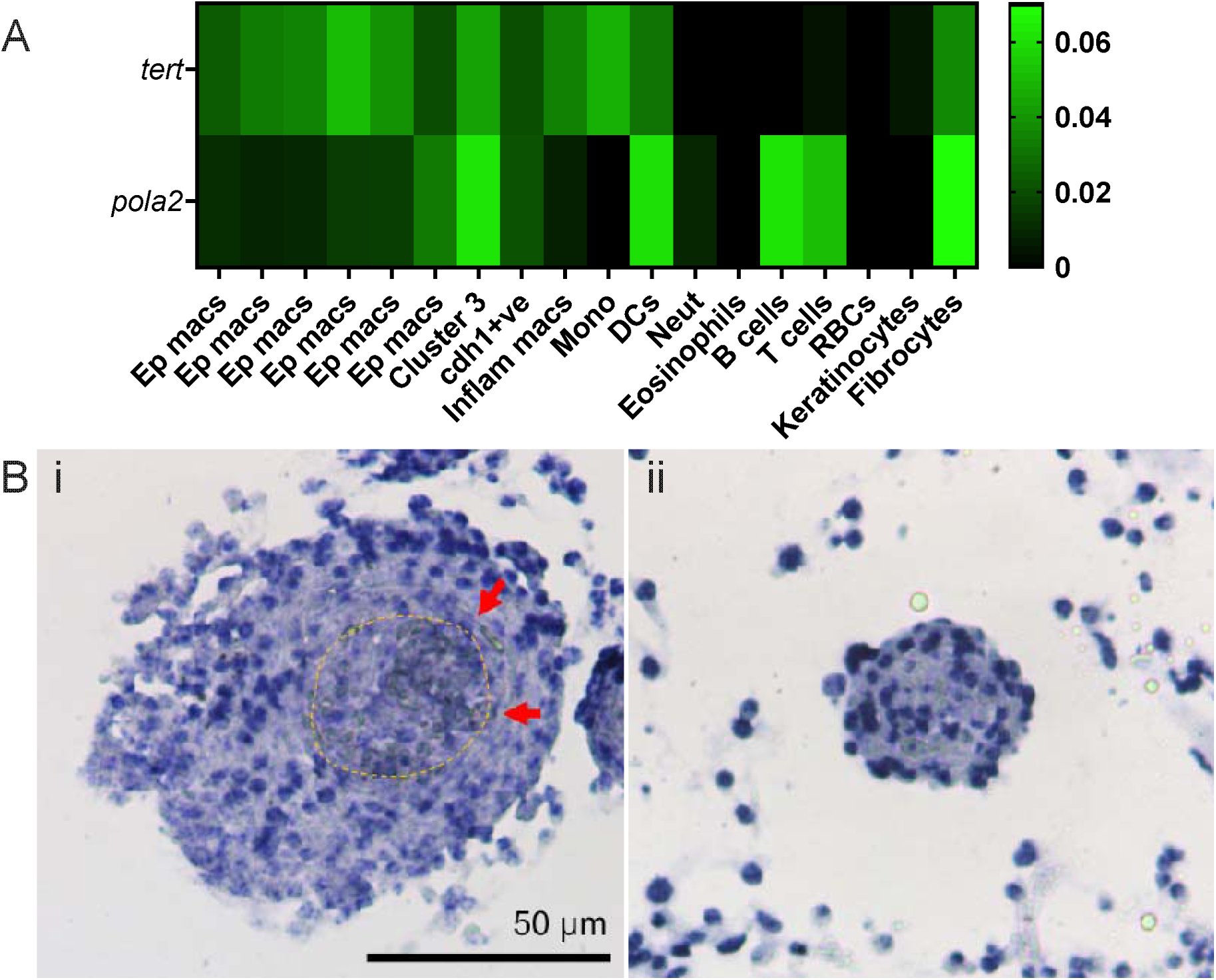
A subset of zebrafish macrophages express *tert* in mycobacterial granulomas. (A) Mean expression value of *tert* and *pola2* across clusters of granuloma cells identified by scRNAseq of adult zebrafish-*M. marinum* granulomas from (*21*). (B) Detection of *tert* expression (blue staining) in (i) necrotic and (ii) cellular non-necrotic adult zebrafish-*M. marinum* granulomas by *in situ* hybridization. Yellow dashed line indicates edge of necrotic core and red arrows indicate epithelioid macrophage layer.

We analyzed the expression of *tert* in adult zebrafish-*M. marinum* granulomas by *in situ* hybridization and found expression in both the flat inner layer of primarily macrophage-derived epithelialized cells and in round cells around the periphery of the granuloma, presumably the proliferating cells identified in the scRNAseq dataset or monocyte-derived cells (Figure 1B). As expected from the heterogenous nature of previously described T-MACs, there were fewer *tert* positive cells detected within granulomas than were seen in previously published images of macrophages visualized with the pan monocyte marker *mpeg1* illustrating expression only in a subset of macrophages (*22*).

### Telomerase deficiency increases the severity of pathogenic mycobacterial infection

We investigated the transcriptional regulation of *tert* following infection by whole embryo RT-qPCR and did not observe changes to *tert* transcription following *M. marinum* infection in developing zebrafish embryos (Supplementary Figure 1A).

We next examined the role of *tert* in the immune response to mycobacterial infection by knocking down expression with a pooled gRNA-CRISPR approach (Supplementary Figure 1B). Knockdown of *tert* increased *M. marinum* burden in zebrafish larvae at 5 days post infection (dpi) (Figure 2A). Our pooled gRNA-CRISPR approach reduced the telomeric repeat amplification protocol (TRAP) activity of whole embryo lysate demonstrating reduced canonical Tert function (Figure 2B). Consistent with telomere shortening being an age-dependent phenotype manifesting over months in zebrafish models, we did not observe any change to telomere length by whole body aggregate qPCR (Supplementary Figure 1C)

**Figure 2:**
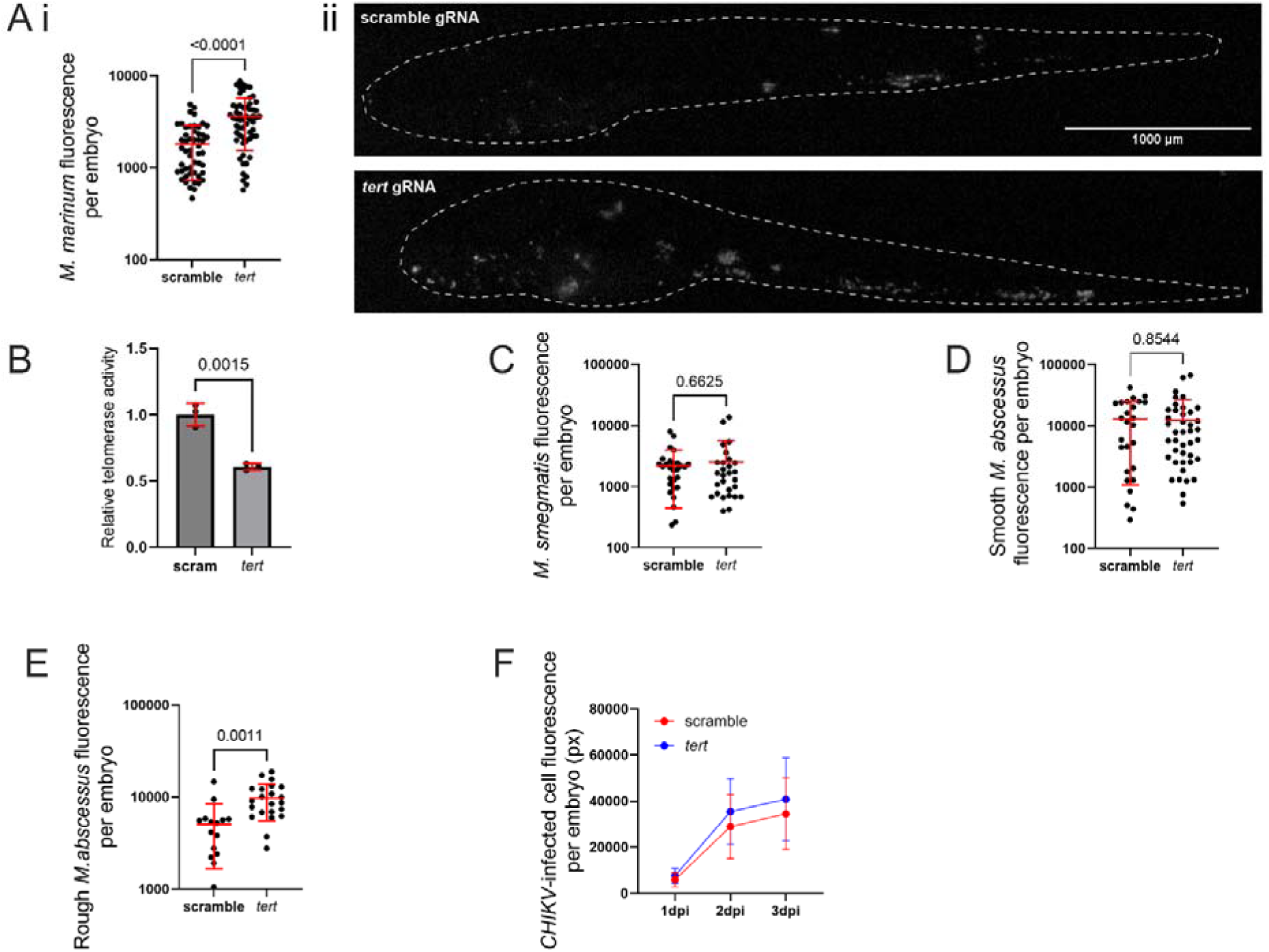
Knockdown of *tert* increases zebrafish susceptibility to pathogenic mycobacterial infection (A) i. Quantification of *M. marinum* burden in 5 dpi *tert* crispant embryos. ii. Representative images of fluorescent *M. marinum* signal in 5 dpi *tert* crispant embryos. Dashed lines indicate outline of embryos. (B) TRAP quantification of relative telomerase activity in 5 dpi *tert* crispant embryos. (C) Quantification of *M. smegmatis* burden in 3 dpi *tert* crispant embryos. (D) Quantification of smooth *M. abscessus* burden in 6 dpi *tert* crispant embryos. (E) Quantification of rough *M. abscessus* burden in 6 dpi *tert* crispant embryos (F) Quantification of CHIKV infected cells in 1-3 dpi *tert* crispant embryos (n= 29 scramble, n= 35 *tert* crispants).

In contrast, infection with two relatively non-pathogenic mycobacteria, *Mycobacterium smegmatis* and smooth colony variant *Mycobacterium abscessus*, was comparable between scramble control and *tert* crispant embryos (Figure 2C and 2D). Infection with the more virulent rough colony variant *M. abscessus* resulted in higher bacterial burden in *tert* crispant embryos (Figure 2E). These data suggest the mycobacterial infection susceptibility is specific to infections capable of forming persistent inflammatory lesions.

We next tested the ability of *tert* crispants to control Chikungunya virus (CHIKV) infection, a well characterized wide host range RNA virus capable of replicating in zebrafish embryos (*23*). Quantification of infected cell area by fluorometry revealed comparable CHIKV infection levels between scramble control and *tert* crispant embryos, demonstrating a specific immune defect to infection by pathogenic bacteria (Figure 2F).

### Loss of the *telomerase RNA component* (*terc*) does not affect mycobacterial infection susceptibility

We complemented our analysis of *tert* by investigating *terc* dynamics and function during *M. marinum* infection in control and *tert*-deficient crispant embryos. We observed a decrease in *terc* levels during mycobacterial infection independent of *tert* status (Supplementary Figure 2A). Knockdown of *terc* using morpholino antisense oligonucleotides reduced the level of *terc* to 24.3 ± 7.1% (n = 4) of control levels (Supplementary Figure 2B) (*17*). However, we did not observe any effect of *terc* knockdown on *M. marinum* infection burden at 5 dpi (Supplementary Figure 2C). Previous reports had demonstrated a role for *terc* in supporting myelopoiesis (*17*), however we did not observe any effect of *terc* knockdown on neutrophil numbers in our infection system (Supplementary Figure 2D). We hypothesized that this was an effect of morpholino dilution allowing a small but sufficient amount of *terc* to escape blockade as the morpholinos were able to reduce early steady state myelopoiesis at 3 days post fertilization (Supplementary Figure 2E), but not late stead state myelopoiesis at 7 days post fertilization equivalent to our 5 dpi timepoint (Supplementary Figure 2F), or total neutrophil numbers (Supplementary Figure 2G).

### Inhibition of telomerase reverse transcriptase increases infection susceptibility and overexpression is protective

We complemented our CRISPR-Cas9 knockdown approach with small molecule TERT inhibitors BIBR1532 and MST-312 (*24, 25*). Treatment of infected embryos increased *M. marinum* burden consistent with our genetic approach data (Figure 3A and 3B).

**Figure 3:**
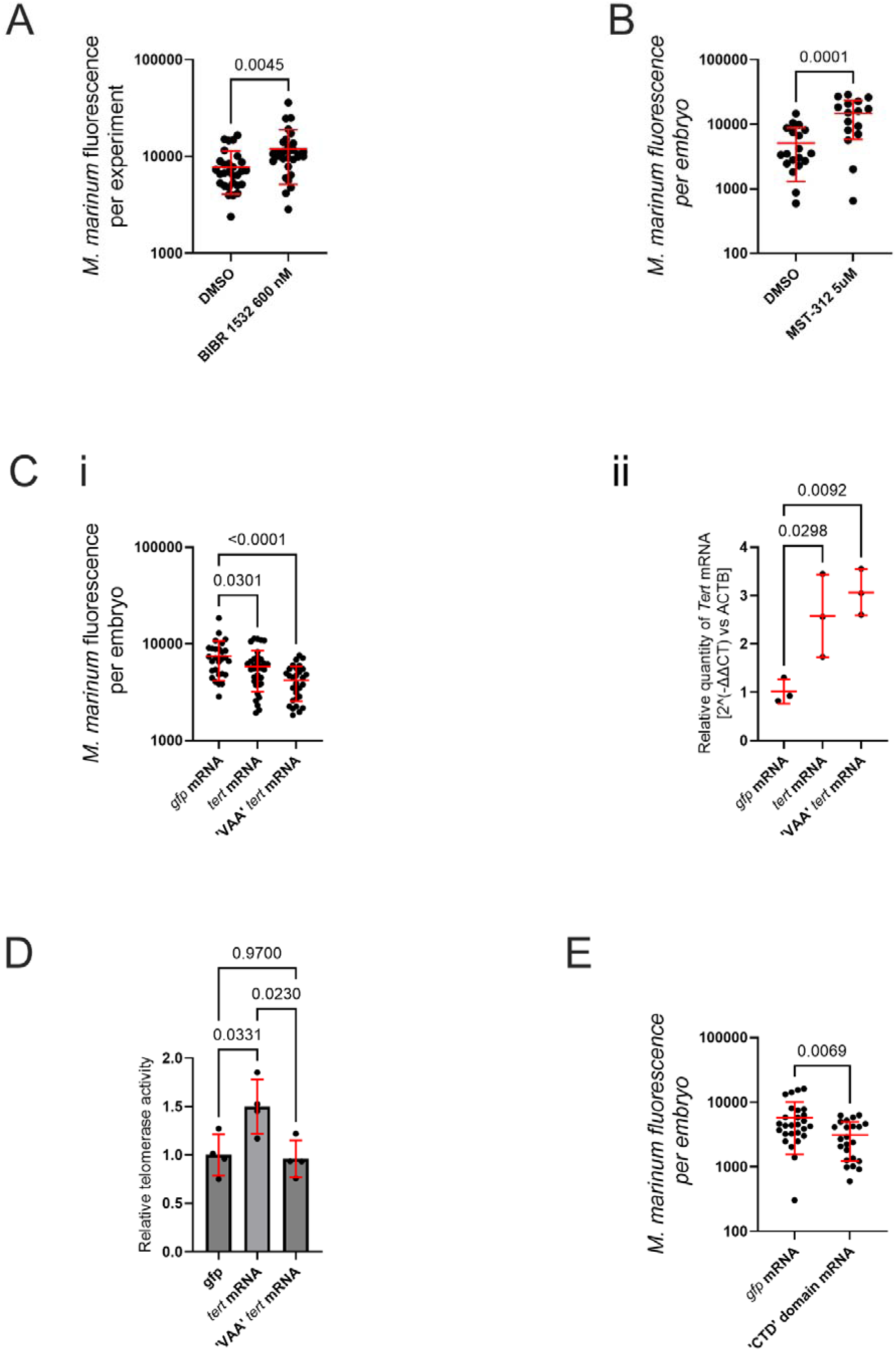
Small molecule and genetic inhibition of Tert increases susceptibility to mycobacterial infection while activation increases resistance to mycobacterial infection. (A) Quantification of *M. marinum* burden in 5 dpi embryos treated with BIBR1532. (B) Quantification of *M. marinum* burden in 5 dpi embryos treated with MST-312. (C) i. Quantification of *M. marinum* burden in 5 dpi embryos injected with control *gfp*, *tert*, or VAA *tert* mRNA. ii. Quantification of *tert* transcript abundance in 1 dpf embryos injected with control *gfp*, *tert*, or VAA *tert* mRNA. (D) TRAP quantification of relative telomerase activity in 5 dpi *tert* and VAA *tert* mRNA-injected embryos. (E) Quantification of *M. marinum* burden in 5 dpi embryos injected with control *gfp* or C-terminal domain (CTD) *tert* mRNA.

Overexpression of *tert* by microinjection with mRNA encoding zebrafish *tert* also reduced *M. marinum* burden (Figure 3C). Interestingly, supplementation of catalytically dead “VAA” *tert* mRNA reduced *M. marinum* burden (Supplementary File 1) (*20*). Analysis of Tert activity by TRAP found increased Tert activity in embryos injected with zebrafish *tert* mRNA but not with “VAA” *tert* mRNA (Figure 3D), suggesting a non-canonical function of Tert could be protective against infection by pathogenic mycobacteria. We further explored the effect of 3′ fragments of *tert* mRNA containing only the C terminus domain (CTD) of the predicted protein (Supplementary File 1), and found a protective effect of overexpression consistent with a non-canonical Tert function (Figure 3E).

### Canonical DNA damage responses do not drive susceptibility to infection in *tert* crispants

Chronic infections reduce LTL and DNA damage-induced interferonopathy plays a critical role in TB pathogenesis (*26, 27*), however previous reports have demonstrated Tert-dependent telomere shortening is only observed in adult zebrafish and not at the embryo stage (*16, 28, 29*). We next investigated the hypothesis that canonical telomere-protective functions of Tert were necessary for resistance to *M. marinum* infection in the context of an increased genotoxic microenvironment caused by infection. Consistent with the presence of an increase in senescent cells, we observed SPiDER-βGal staining around *M. marinum* granulomas demonstrating the presence of the senescent cell marker beta galactosidase activity (Figure 4A).

**Figure 4:**
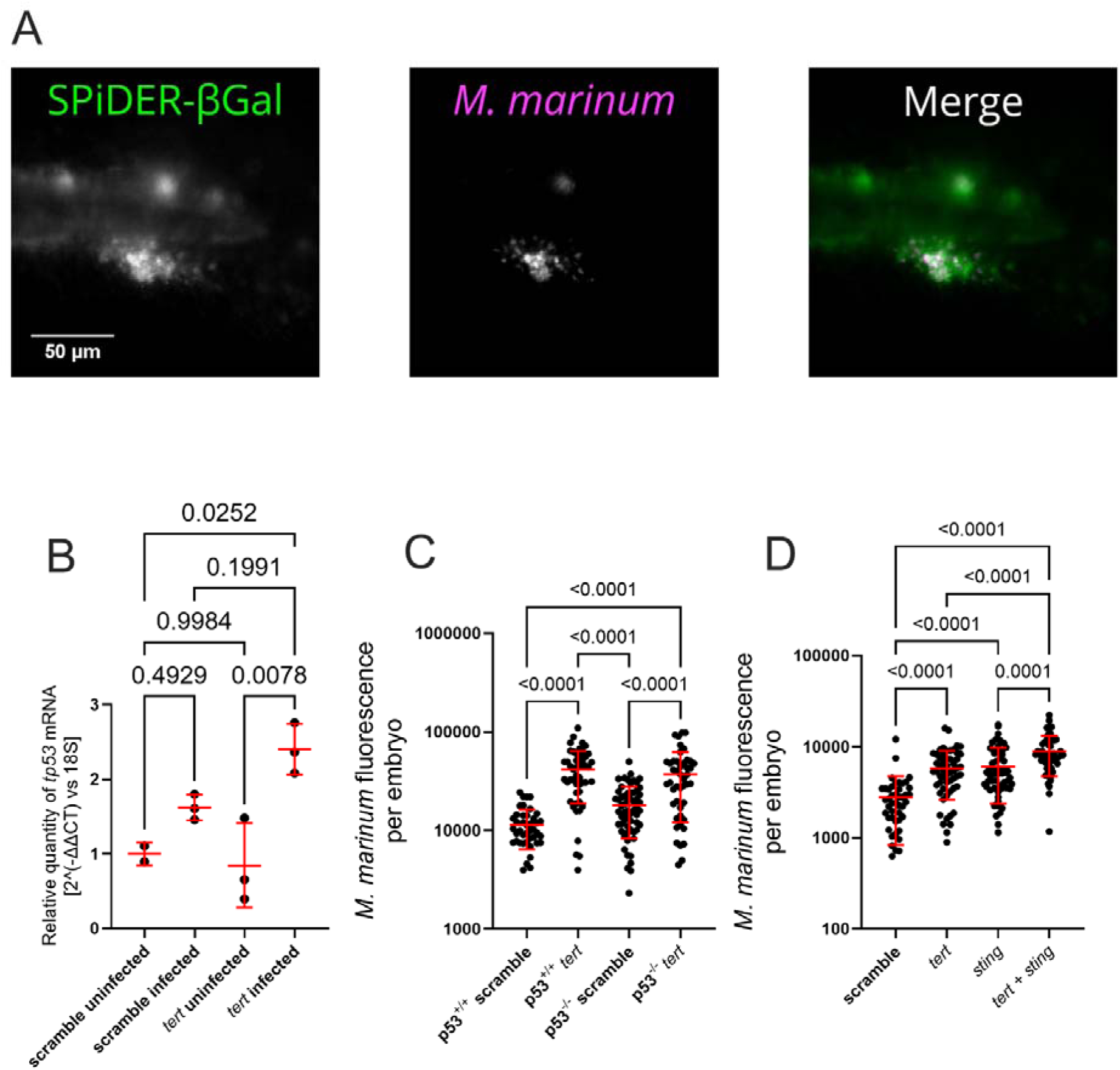
Increased susceptibility to infection in *tert* crispants is not driven by DNA damage response pathways. (A) Fluorescent SPiDER-βGal staining (green) around *M. marinum* granuloma (magenta) in a 5 dpi embryo. (B) Quantification of *tp53* transcript abundance in 5 dpi *tert* crispant embryos infected with *M. marinum*. (C) Quantification of *M. marinum* burden in 5 dpi *tp53* mutant embryos. (D) Quantification of *M. marinum* burden in 5 dpi *sting* crispant embryos.

Damage to chromosomal DNA has been demonstrated to activated Tp53-mediated growth restriction and Sting-mediated inflammation to accelerate aging phenotypes in Tert deficient adult zebrafish (*30, 31*). We hypothesized arrest of host immune cell proliferation and Sting-mediated production of type I interferons would be detrimental to control of *M. marinum* infection. Expression of the DNA damage response gene *tp53* was increased in infected *tert* crispant embryos compared to uninfected *tert* crispants indicating a small global increase in DNA damage in the infection-susceptible *tert* crispants (Figure 4B).

However, functional examination of the DNA damage response in *tp53* mutants and *sting* crispants failed to rescue the *tert* knockdown-induced susceptibility to *M. marinum* infection in our embryo infection model suggesting a non-canonical role for Tert in our system(Figure 4C and 4D, Supplementary Figure 3).

### Tert supports infection-induced myelopoiesis

Zebrafish *tert* and *terc* are known to be necessary for developmental myelopoiesis (*16, 17, 32, 33*), and *M. marinum* infection control requires replenishment of the myeloid cell compartment (*34*). Conversely, infection with smooth morphotype *M. abscessus* or *M. smegmatis*, two infections that were not exacerbated by Tert depletion, did not induce myelopoiesis in zebrafish embryos (Supplementary Figure 4A and 4B). Furthermore, we did not observe a specific leukocytic response to the sites of CHIKV replication correlating Tert-dependent immunity with severe infections that drive infection-induced myelopoiesis (Supplementary Figure 4C).

We next analyzed infection-induced myelopoiesis as a potential mechanism of susceptibility in *tert* crispants using *Tg(lyzc:GFP)^nz117^* and *Tg(lyzc:dsred)^nz50^* lines, where neutrophils are fluorescently labelled (*35*), and the *TgBAC(mpeg1.1:EGFP)^vcc7^* line where macrophages are fluorescently labelled (*36*). Caudal hematopoietic tissue-resident neutrophil and macrophages were reduced in *tert* crispants late in infection (Figure 5A and 5B).

**Figure 5:**
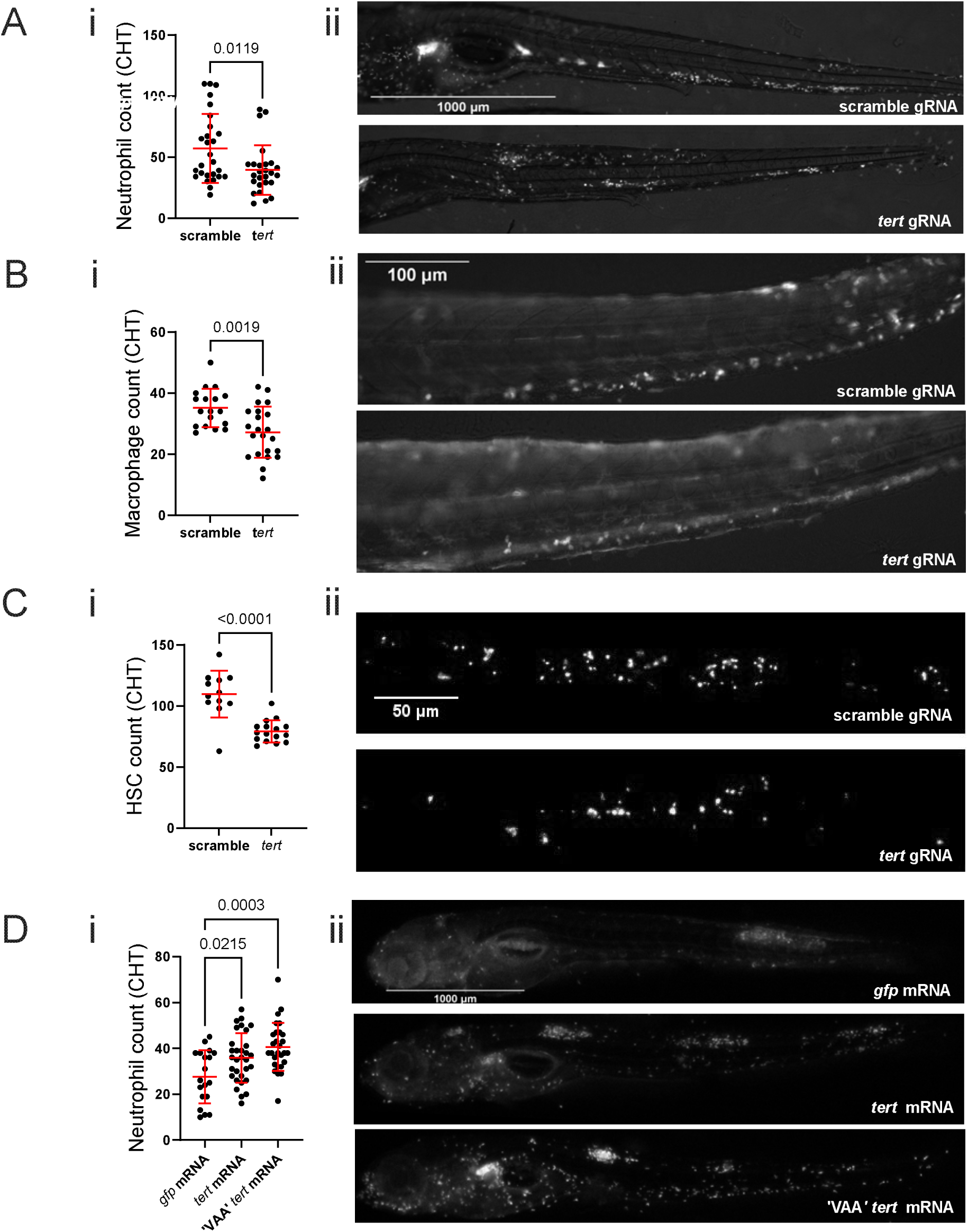
Expression of *tert* correlates with infection demand-driven leukocyte abundance. (A) i. Quantification of CHT-resident neutrophils in 5 dpi *Tg(lyzC:dsred) tert* crispant embryos infected with *M. marinum*. ii. Representative images of fluorescent neutrophils. Bracket demarcates CHT region used for counting. (B) i. Quantification of CHT-resident macrophages in 5 dpi *TgBAC(mpeg1.1:gfp) tert* crispant embryos infected with *M. marinum*. ii. Representative images of fluorescent macrophages in the CHT region. (C) i. Quantification of CHT-resident HSCs in 5 dpi *Tg(Mmu.Runx1:NLS-mCherry)* embryos infected with *M. marinum*. ii. Representative images of fluorescent HSCs in the CHT region. (D) i. Quantification of CHT-resident neutrophils in 5 dpi *Tg(lyzC:dsred)* embryos injected with control eGFP, *tert*, or VAA *tert* mRNA and infected with *M. marinum*. ii. Representative images of fluorescent neutrophils.

To establish if the loss of *lyzc* and *mpeg1.1* positive cells was due to reduced haematopoietic stem cell (HSC) numbers or a lack of differentiation of myeloid cells, we enumerated *Tg(Mmu.Runx1:NLS-mCherry)^cz2010^* positive HSCs (Figure 5C) (*37*). Tert crispants exhibited a decline in HSC counts at 5 dpi, corresponding with the observed loss of myeloid cells.

Genetic supplementation of endogenous *tert* by microinjection with mRNA encoding full length and catalytically dead zebrafish *tert* increased neutrophil numbers late in infection (Figure 5D). This increase in leukocyte number corresponded with protection from infection (Figure 3C) placing Tert in the control pathogenic mycobacterial infection via supporting infection-induced myelopoiesis.

## Discussion

Our study used depletion of zebrafish telomerase reverse transcriptase to find a novel role for telomerase in the control of mycobacterial infection via supporting infection demand-driven myelopoiesis. These data provide a mechanism that may contribute to the increasing susceptibility to TB during ageing and provide a complementary explanation for the association of short LTL with infection susceptibility. The data further suggest supplementation of telomerase or myelopoietic factors could be used to bolster anti-TB immunity in the elderly (*34*).

While disruption of telomerase activity is associated with canonical telomere-shortening, our use of the VAA active site mutant *tert* and CTD *tert* mRNA to curtail *M. marinum* infection points towards a non-canonical function of Tert in infection demand-driven hematopoiesis. We note that although BIBR1532 binds to *Tribolium castaneum* TERT near the nucleotide binding domain, although it is unclear if this causes loss of non-canonical functions in addition the well-documented loss of canonical function (*38*). Furthermore, the mechanism of action for MST-312 remains unknown. We have previously reported a non-canonical role of zebrafish Tert in supporting gut macrophage proliferation in adult animals (*19*), an organ with constant turnover and immune stimulation, and consistent with the defect in infection demand-driven hematopoiesis found in this study. Non-canonical telomerase, non-coding RNA component (TERC), and shelterin functions span a wide range of cellular activities, while there are similarities in the DNA-based themes of these functions there is also clear mechanistic independence of their non-canonical activities (*39, 40*).

Disseminated *M. tuberculosis* drives epigenetic reprogramming of HSCs to impair the myelopoietic response of infection (*41*). Our zebrafish-*M. marinum* infection model data adds to this concept by demonstrating loss of HSCs sensitized by the loss of host telomerase, suggesting TB infection may accelerate the onset of clonal hematopoiesis in the elderly. Our finding that the premature aging model of telomerase deficiency is a susceptibility factor for pathogenic mycobacterial infection ties together with prior human TB and guinea pig-*M. tuberculosis* infection model data finding *M. tuberculosis* infection prematurely ages the host (*4*). Together, these data suggest a vicous cycle whereby TB ages the host triggering a premature decline of hematopoietic function and further reducing the ability of the host to resist infection.

The loss of HSCs in our chronic mycobacterial infection model suggests TERT may safeguard HSCs from infection-induced damage in addition to performing previously described roles as an inflammatory transcription factor (*42*). Although we did not observe bulk effects of *tert* loss on telomere length in our embryos, we must include the possibility that actively dividing HSCs are more susceptible to loss of canonical telomerase activity causing them to enter premature replicative senescence in chronic infection. We have recently identified an immune cell-autonomous role of Tert in supporting STING-TBK1 signaling (*20*), that while conventionally considered to reduce pathogenic Type I Interferon in chronic murine *M. tuberculosis* infection, may reduce early immune activation in non-murine mycobacterial infection systems (*43*).Further phenotyping of the HSC and macrophage compartment of TERT mutant animals in the context of infection is required to determine the range of roles played by telomerase in protecting HSCs and macrophage-mediated immune defense.

Multiple studies have linked short telomere length as a proxy for reduced telomerase expression to viral infection susceptibility in species as diverse as humans and rock oysters (*5, 10–12, 44*), however little is known about the mechanistic basis for this association. The only prior mechanistic data connecting telomerase to infection susceptibility is the cell-intrinsic hijacking of host telomerase to support viral replication during lytic human cytomegalovirus infection (*45*). Our bacterial infection experiments add a new dimension to the literature documenting interactions between telomerase and infection across host and pathogen species. We hypothesize the relative contributions of HSC and T-MAC telomerase expression will manifest in a disease-specific manner and encourage more investigation of telomerase function in diverse disease settings.

## Methods

### Animal husbandry

Zebrafish experiments were approved under the A*STAR IACUC protocols 211667 and 221694; Shanghai Public Health and Clinical Center, Fudan University GB/T 35823-2018 and 2022-A002-01; UK Home Office Project License PPL PP5900632. Breeding adults were housed under 14 hour light / 10 hour dark cycles in a 28°C recirculating aquarium. Eggs were produced by natural spawning and embryos were raised in E3 media at 28°C and supplemented with PTU from 24 hours post fertilization.

### Bacterial culture

*M. abscessus, M. marinum,* and *M. smegmatis* were cultured in 7H9 or on 7H10 supplemented with OADC and hygromycin to select for pTEC fluorescent protein plasmids (L. Ramakrishnan, plasmids are available through Addgene https://www.addgene.org/Lalita_Ramakrishnan/). Single cell suspensions of mycobacteria were prepared and frozen at −70°C in aliquots for individual experiments as previously described (*46*).

### Viral culture

Chikungunya virus (CHIKV) virus was produced from an infectious clone plasmid containing a duplicated subgenomic promoter driving ZsGreen (*Anthozoa* reef coral fluorescent protein) expression between the non-structural and structural genes. The viral genome was flanked by a CMV and a HDV ribozyme to launch transcription. BHK cells (RRID: CVCL_1914) were transfected with 500 ng of plasmid per well in a 6 well plate using lipofectamine 2000 (Thermofisher). At 3 days post transfection when CPE reaches approximately 40%, supernatant was cleared of cell debris and used to expand CHIKV culture on VeroE6 cells (RRID: CVCL_0574) for 44 hours. Cleared supernatant was tittered by standard plaque assay on VeroE6 cells. VeroE6 and BHK were grown in DMEM high glutamine (Gibco) with 10% FBS (Hyclone).

### Infection with fluorescent pathogens and quantification of infection phenotypes by imaging

Zebrafish embryos were infected by microinjection with approximately 200 CFU *M. marinum*, 1000 CFU DESX1 *M. marinum* or *M. abscessus*, or 50 PFU CHIKV. Primary infection was performed by injection into the caudal vein or neural tube above the yolk sac extension at 2 dpf. Infection was assayed by stereomicroscopy of entire immobilized zebrafish embryos on a Nikon SMZ25 stereoscope or an inverted Nikon Eclipse Ti2 at 5 dpi unless otherwise mentioned. Image analysis was performed in ImageJ/FIJI with bacterial fluorescent pixel count (FPC) carried out as previously described (*47*).

HSC counts were carried out on a Zeiss Celldiscoverer 7 (CD7) widefield microscope equipped with a 20x/0.95 plan-apochromat objective and 0.5x tube lens to acquire a 3 Z-stacks every 5 μm.

### Protein extraction and quantification

8-15 zebrafish embryos were pooled and manually homogenised with pestle (Sigma BAF650000002-100EA) for protein extraction with 30μL of Pierce IP Lysis Buffer (Thermo Fisher 87787) with 1x Halt Protease and Phosphatase Inhibitor Cocktail (Thermo Fisher 78440) on ice for 30 minutes. Extract was centrifuged at 16,000g for 30 minutes at 4°C. Supernatant were collected and diluted 20 folds before protein concentration was quantified using Pierce™ BCA Protein Assay Kits (Thermo Fisher 23225) in a Tecan plate reader at 562nm. Samples were extracted as pooled triplicates.

Quantitative telomeric repeat amplification protocolThe standard curve was constructed using 1000ng, 200ng, 40ng, 8ng, 1.6ng of protein per experiment and 10^[(Ct^ ^sample-Yint)/slope]^ was used to determine the relative telomerase activity of samples. Per 25μL reaction, 8.5μL of nuclease free water, 12.5μL of 2x PowerUp SYBR Green Master Mix (Thermo Fisher A25742), 1μL of 500ng/μL of TS and ACX primer and 2μL of 500ng of protein extract was used. CFX96 Touch Real-Time PCR Detection System was used to run the following protocol: 30°C for 30 minutes, 95°C for 10 minutes, 40 cycles of 95°C for 15 seconds and 60°C for 1 minute.

### *In situ* hybridization

DIG-labeled RNA antisense probes were synthesized by *in vitro* transcription with DIG-labeled mix (Roche 11277073910) and the MEGA shortscript T7 Transcription kit (Thermo Fisher AM1354) from *tert* cDNA cloned into a pBluescript SK vector (primers referred to as ISH-tert in Table 1). Probe preparation and *in situ* hybridization of paraffin sections were carried out as described (*22*).

**Table 1:**
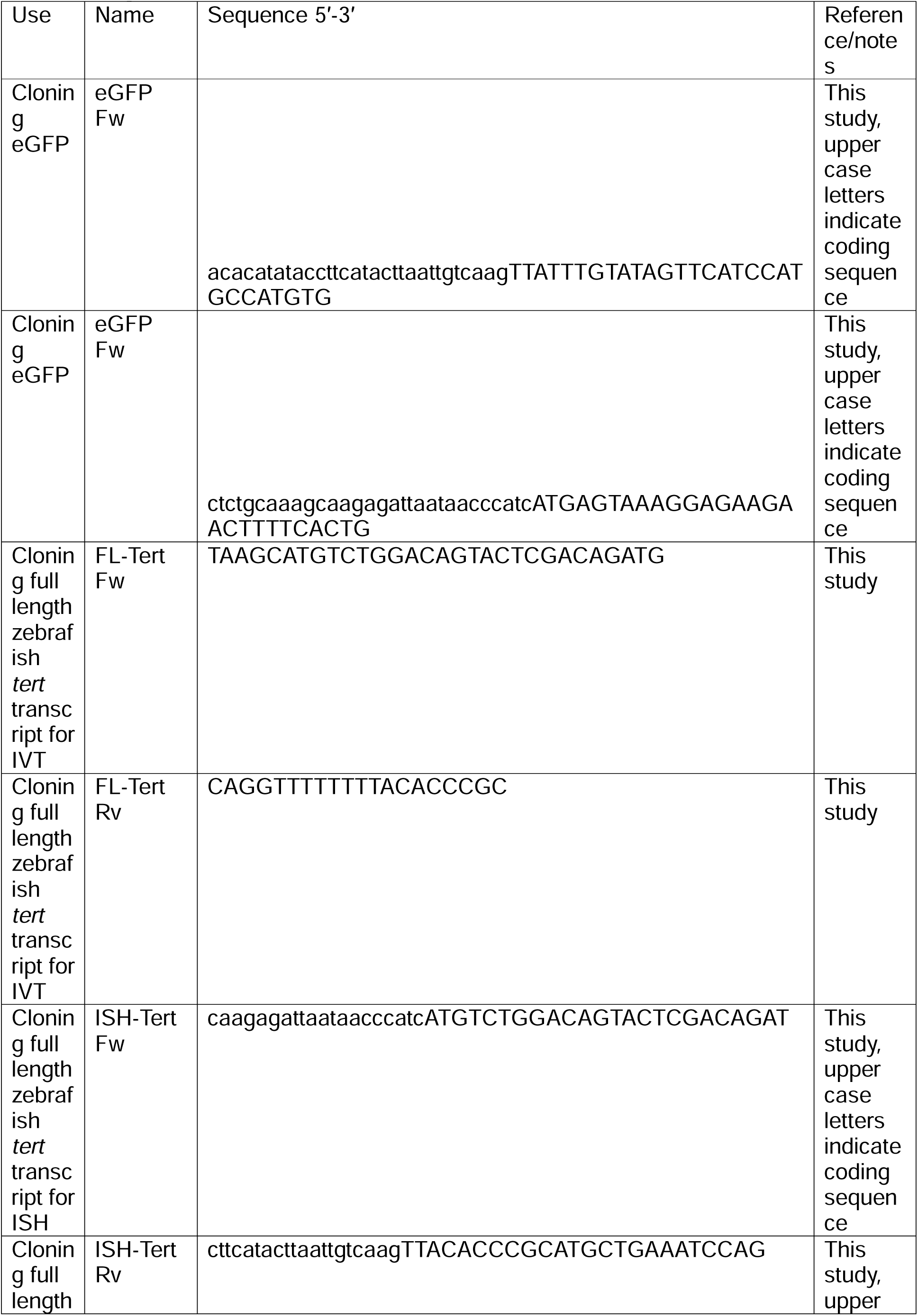
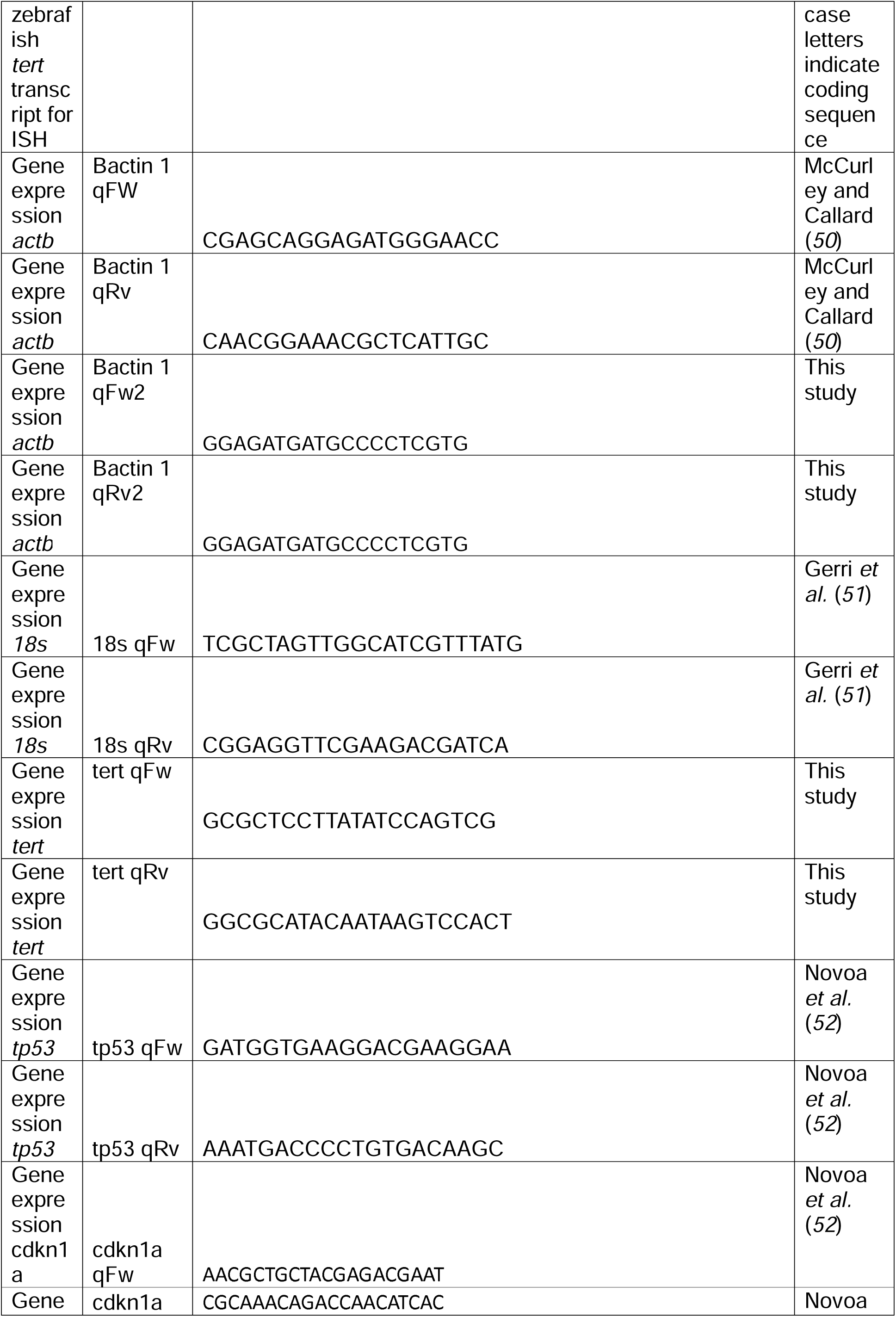
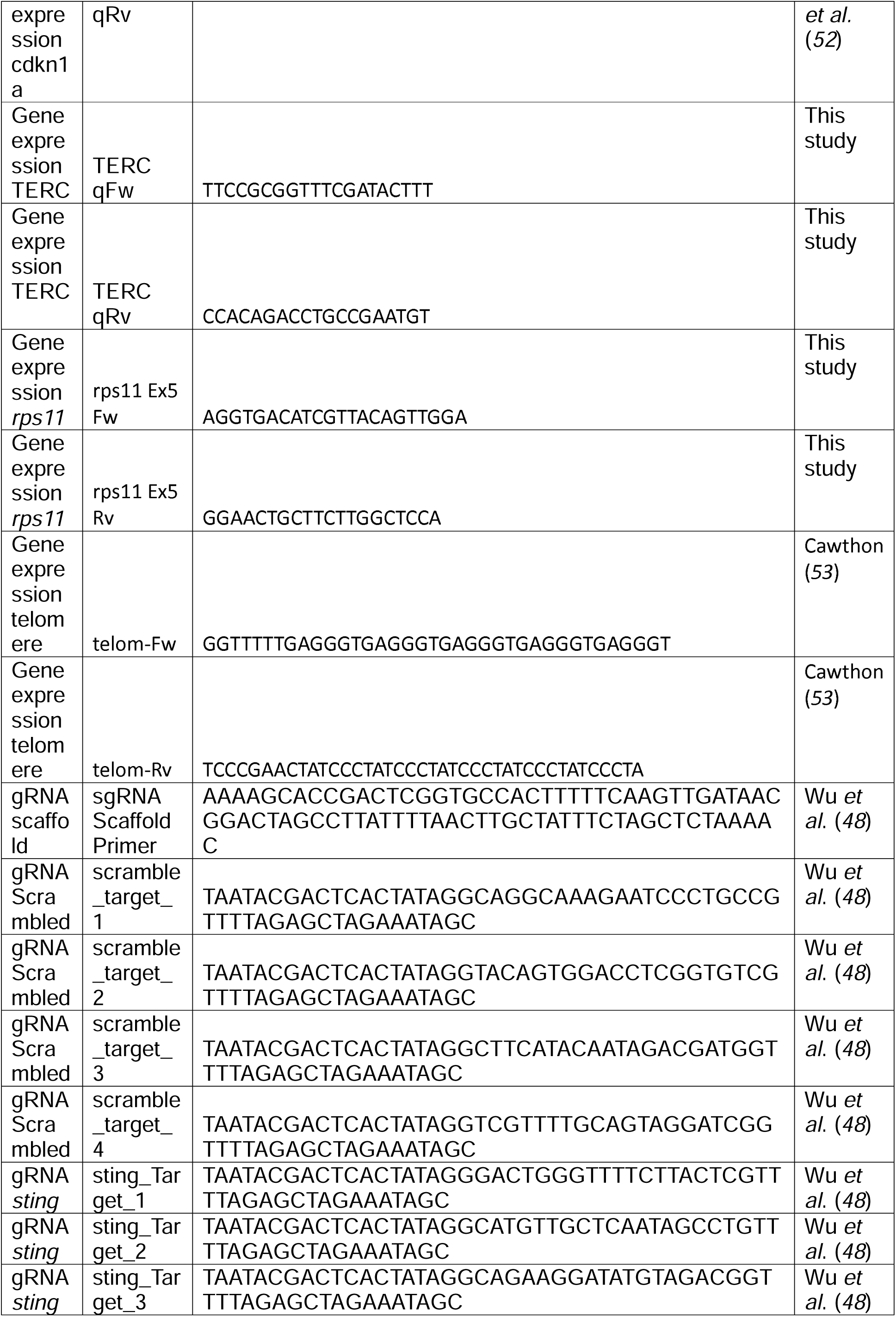
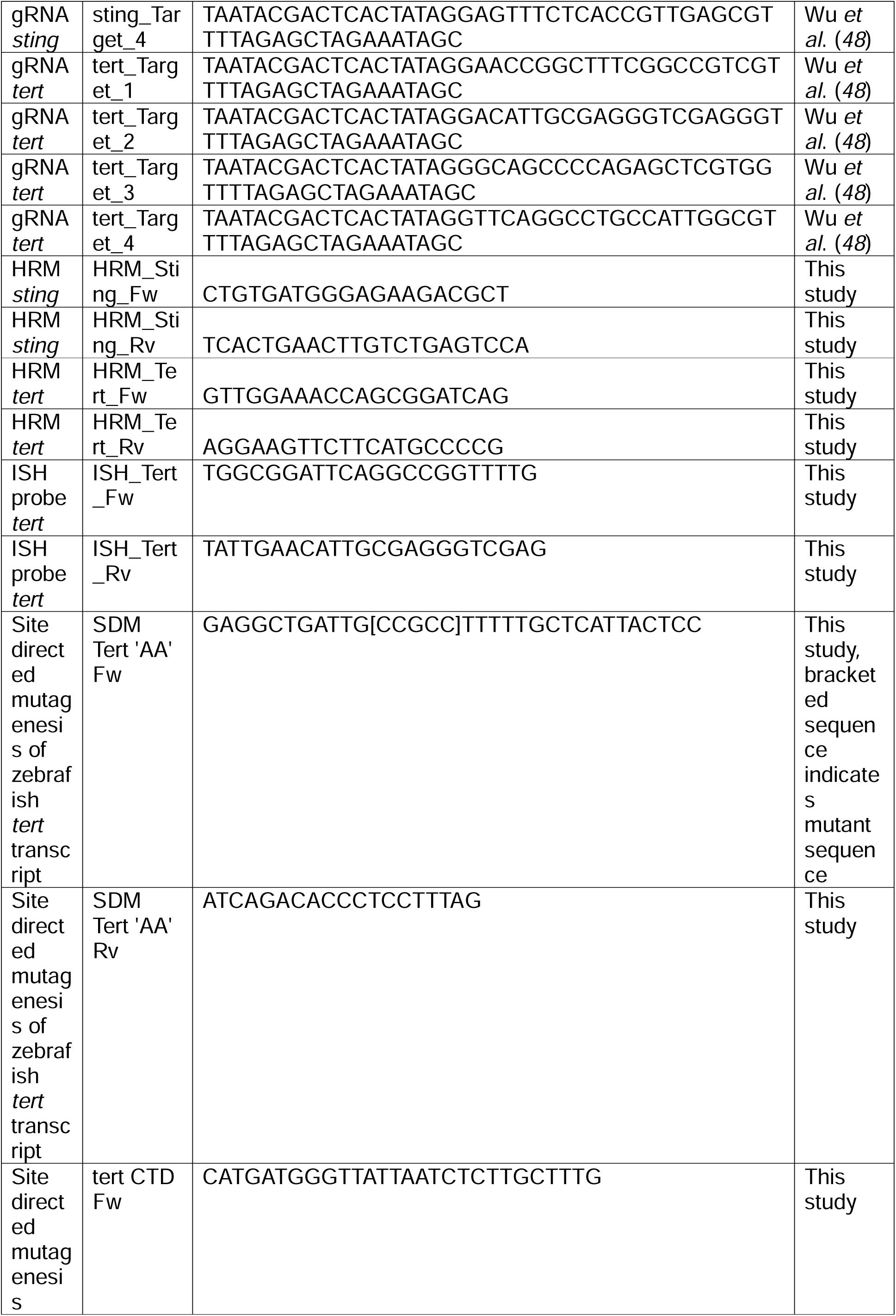
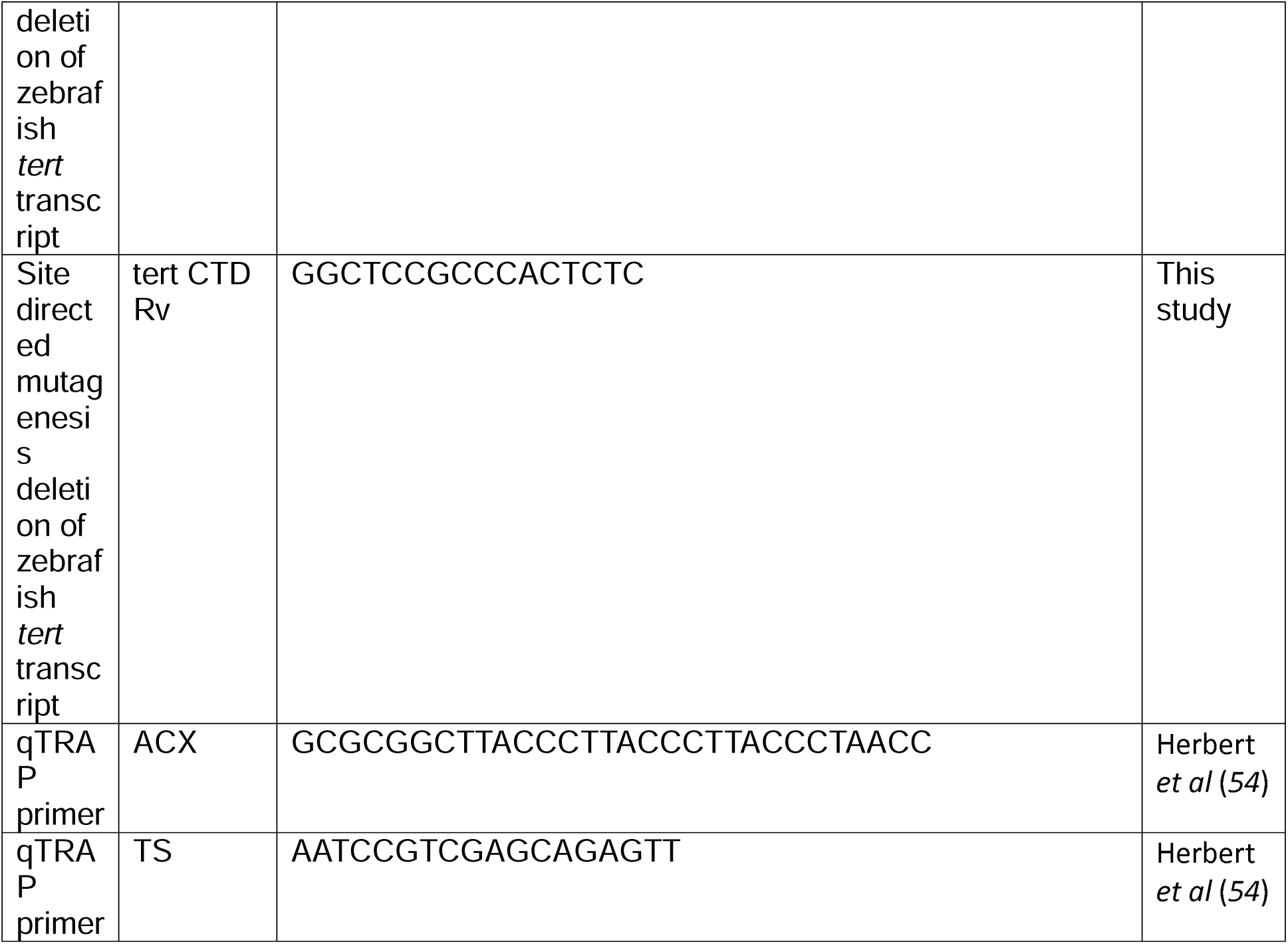
Oligonucleotides.

### CRISPR-Cas9 mutagenesis

Gene-specific oligos for *in vitro* transcription of gRNAs were designed with IDT or selected from Wu *et al*. (*48*). HiScribe T7 High Yield RNA Synthesis Kit (NEB E2040) was used for *in vitro* transcription and Alt-R CRISPR-Cas9 crRNA was purchased commercially (IDT). Alt-R S.p. Cas9 Nuclease V3 (IDT 1081059) was combined with gRNA/crRNA and injected into the yolk of 1-2 cell stage zebrafish embryos.

### Small molecule treatment

BIBR1532 (Selleckchem S1186)and MST-312 (Sigma M3949) were dissolved in DMSO. Small molecules were added to the media of infected embryos at final concentrations of 600 nM BIBR1532 and 2.5 μM MST-312. Small molecules were refreshed in complete media changes every second day.

### *In vitro* mRNA synthesis and embryo injection

A full length *tert* cDNA (referred to as FL-tert in Table 1) was cloned into pGEMTeasy using Q5 polymerase (New England Biolabs). This was subcloned into a pSP64 Poly(A) plasmid (Promega P1241) using NEBuilder HiFi DNA Assembly (NEB E2621). Site-directed mutagenesis was carried out with the Q5 Site-Directed Mutagenesis Kit (NEB E0554) to mutate the ATGAT bases to CCGCC resulting in a predicted change in aspartate to alanine recreating the VAA catalytically dead TERT mutation (*49*). *In vitro* transcription of capped mRNA was performed with mMESSAGE mMACHINE Sp6 Transcription Kit (Invitrogen AM1340). tert CTD primers (Refer to Table 1) was used to generate a truncated C terminal domain (CTD) *tert* construct from full length *tert* cDNA with Q5 polymerase and KLD enzyme master mix (NEB M0554). *In vitro* transcription of capped mRNA was performed with mMESSAGE mMACHINE Sp6 Transcription Kit (Invitrogen AM1340). 300 pg of mRNA, in 1.5 nL volume, was injected into the yolk of 1-4 cell stage zebrafish embryos.

### DNA extraction and telomere length analysis

7-10 embryos were pooled for gDNA extraction using Monarch Spin gDNA Extraction Kit (NEB T3010L) as per manufacturer’s protocol with minor adjustments, 2 μL of proteinase K and RNase A was used instead. Samples were extracted as pooled triplicates. Monarch Spin PCR & DNA Cleanup Kit (5 μg) (NEB T1130L) was used to purify and concentrate gDNA.

30 ng of gDNA was used per 10 μL reaction with *rps11* as the single copy gene housekeeper. Per 10 μL reaction, 3.2 μL of nuclease free water, 5 μL of 2x PowerUp SYBR Green Master Mix (Thermo Fisher A25742), 0.4 μL of 10 μM forward and reverse primers and 1 μL of gDNA was used. 2−ΔΔ.Ct was used to analysis relative telomere length.

### DNA damage assays

SPiDER-βGal staining was carried out with the Cellular Senescence Detection Kit - SPiDER-βGal (Dojindo SG03-10) as per manufacturer’s instructions on fixed zebrafish embryos equilibrated to McIlvain buffer pH6 and stained for 30 minutes at 37°C.

### RNA extraction

7-25 embryos were pooled for RNA extraction using TRIzol (Thermo Fisher 15596018) or NucleoSpin RNA kit (Macherey-Nagel 740955.250) as per manufacturer’s protocol. Samples were extracted as pooled triplicates.

### Gene expression analysis

Quantitative PCR was performed with reverse transcription using the High Capacity cDNA Kit (Applied Biosystems 4368813) or SensiFAST cDNA Synthesis Kit (BIO-65054) and PCR amplification with PowerUp SYBR Green Master Mix (Thermo Fisher A25742).

### Statistics

Analyses of quantitative data were carried out with Graphpad Prism using T-tests for pairwise comparisons or ANOVA for multiple comparisons. All data are representative of at least 3 biological replicates unless otherwise stated in the captions. Error bars on graphs represent standard deviation.

## Acknowledgements

This study was funded by the Singapore Ministry of Health’s National Medical Research Council under its individual research grant scheme (OFIRG22jul-0081) to S.H.O., European Research Council Consolidator Grant (772853 - ENTRAPMENT) and Wellcome Discovery Award (226644/Z/22/Z) to S.M., the National Natural Science Foundation of China (82372263 to B.Y; 82402628 to G.L).

The authors thank A*STAR IMCB Zebrafish Core Facility discussions and the LSHTM Biological Services Facility for expert animal husbandry; Dr Sudipto Roy for providing the *tp53* mutant zebrafish line; Drs Miguel Ferreira, Oliver Dreesen, Dennis Kappei, and Maya Jeitany for helpful telomerase discussions; Mostowy lab members for helpful discussions; Ms Terumi Nakashima, and Drs Caroline Wee, Shifeng Xue, and Zhipeng Tai for technical assistance.

## Data Availability Statement

The data that support the findings of this study are available from the corresponding author upon reasonable request.

## Author’s Contributions

Darryl JY Han: experimentation, designed experiments, wrote draft David M Costa: experimentation, designed experiments, approved draft Geyang Luo: experimentation, approved draft Kathryn Wright: experimentation, approved draft Manitosh Pandey: experimentation, approved draft Denise Wee: experimentation, approved draft Yao Chen: experimentation, approved draft

Semih Can Akincilar: provided tools, designed experiments, approved draft Vinay Tergaonkar: provided tools, approved draft

Rajkumar Dorajoo: experimentation, approved draft Guillaume Carissimo: provided tools, approved draft Amit Singhal: experimentation, approved draft Philip M Elks: experimentation, approved draft Pamela S Ellis: provided tools, approved draft

Catarina M Henriques: provided tools, designed experiments, approved draft Bo Yan: experimentation, approved draft

Serge Mostowy: experimentation, approved draft

Stefan H Oehlers: conceived study, experimentation, wrote draft

---

**Supplementary Figure 1:**
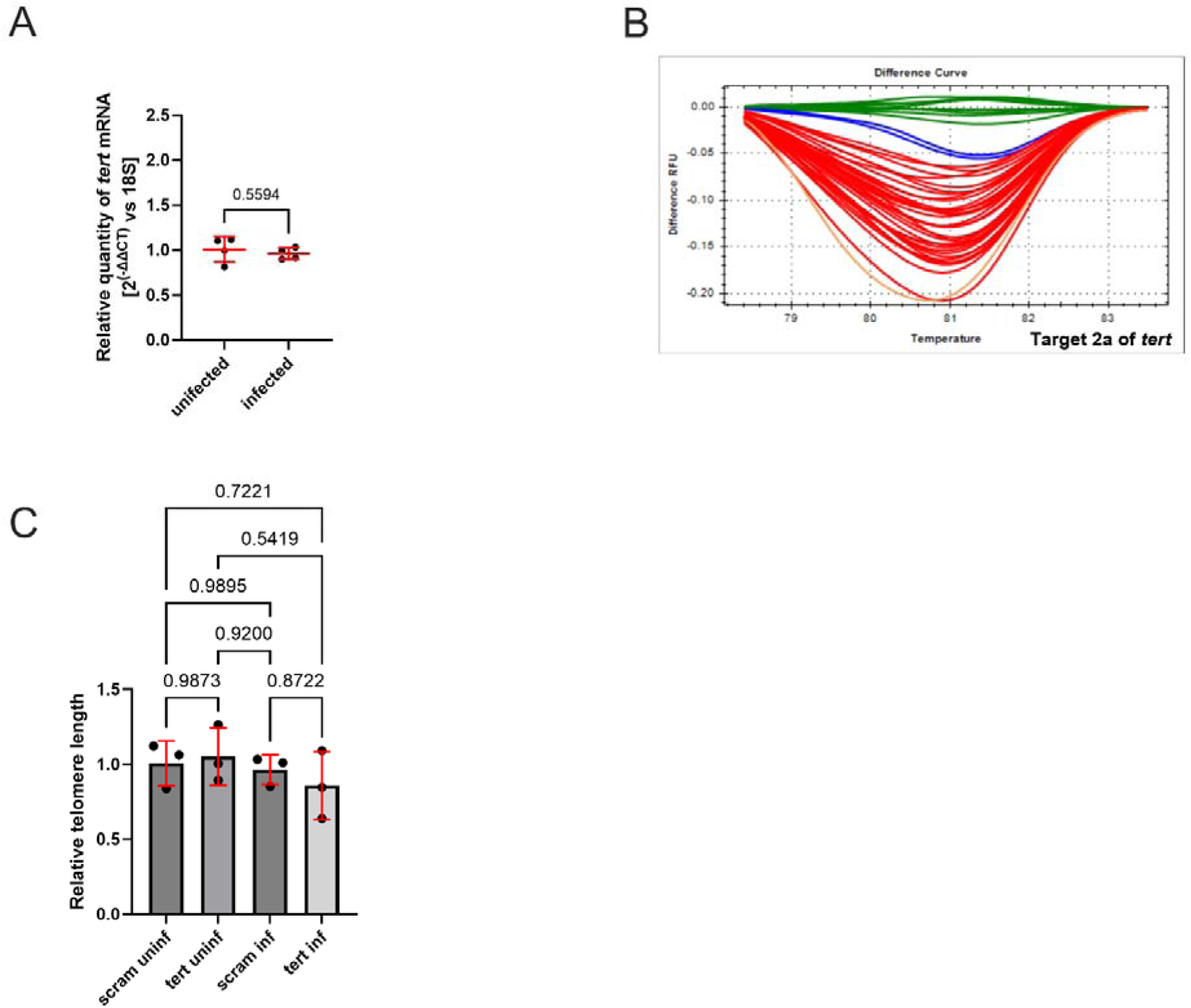
Quantification of *tert* and editing efficiency of pooled gRNA-Cas9 editing of *tert*. (A) Quantification of *tert* in 5 dpi embryos infected with *M. marinum*. (B) High resolution melt analysis of *tert* gRNA2 target site, n=35, 100% efficiency, control clusters: Green & Blue. (C) Quantification of telomere length by qPCR from homogenates of 5 dpi embryos infected with *M. marinum*.

**Supplementary Figure 2:**
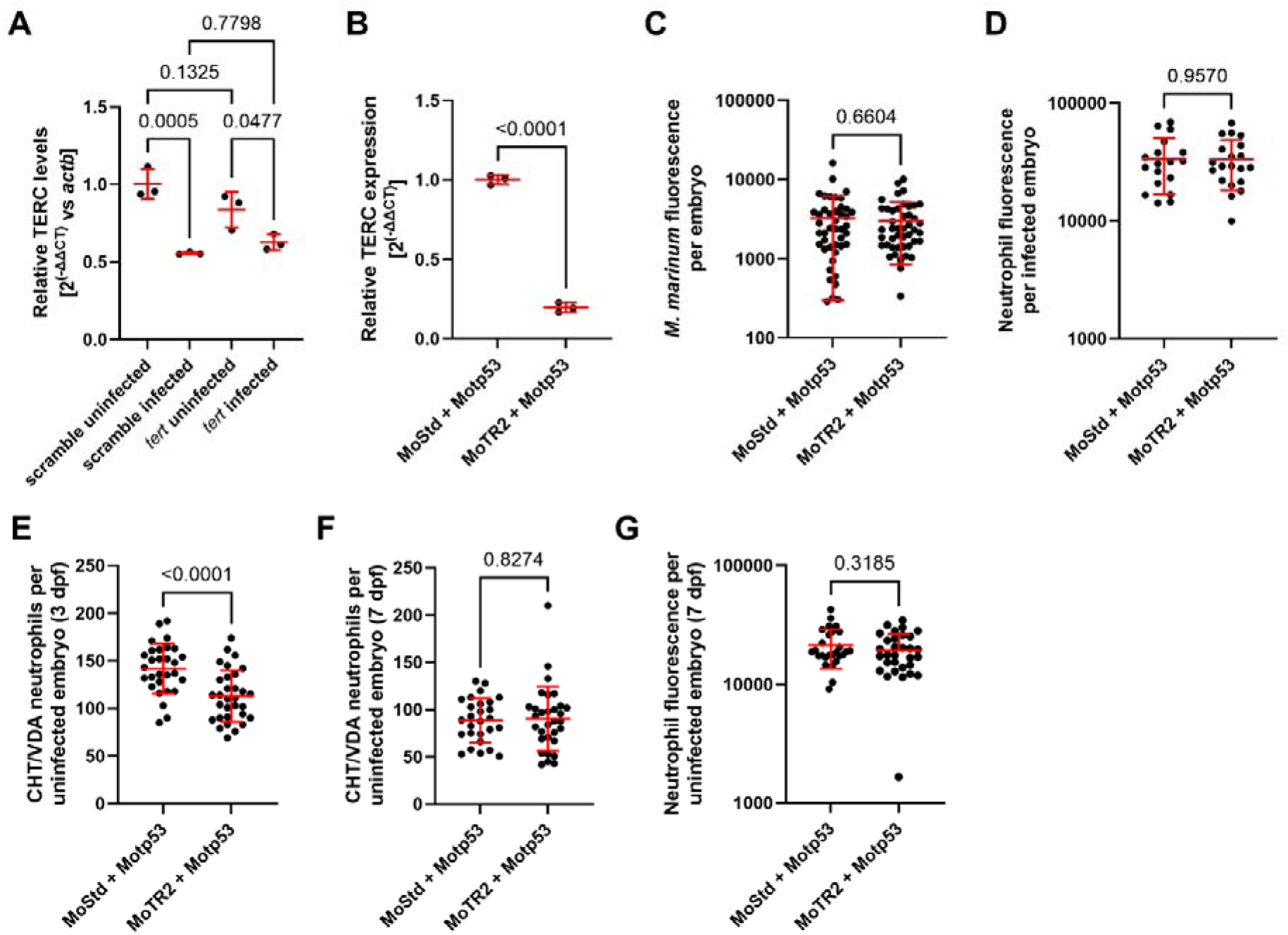
Infection-induced reduction in TERC does not correlate with mycobacterial infection susceptibility in zebrafish embryos (A) Quantification of TERC by qRT-PCR in 5 dpi *tert* crispant embryos infected with *M. marinum*. (B) Quantification of TERC by qRT-PCR in 5 dpi morpholino-injected embryos infected with *M. marinum*. (C) Quantification of *M. marinum* burden in 5 dpi morpholino-injected embryos infected with *M. marinum*. (D) Quantification of total neutrophils in 5 dpi morpholino-injected embryos infected with *M. marinum*. (E) Quantification of CHT region neutrophils in 3 days post fertilization morpholino-injected embryos. (F) Quantification of CHT region neutrophils in 7 days post fertilization morpholino-injected embryos. (G) Quantification of total neutrophils in 7 days post fertilization morpholino-injected embryos.

**Supplementary Figure 3:**
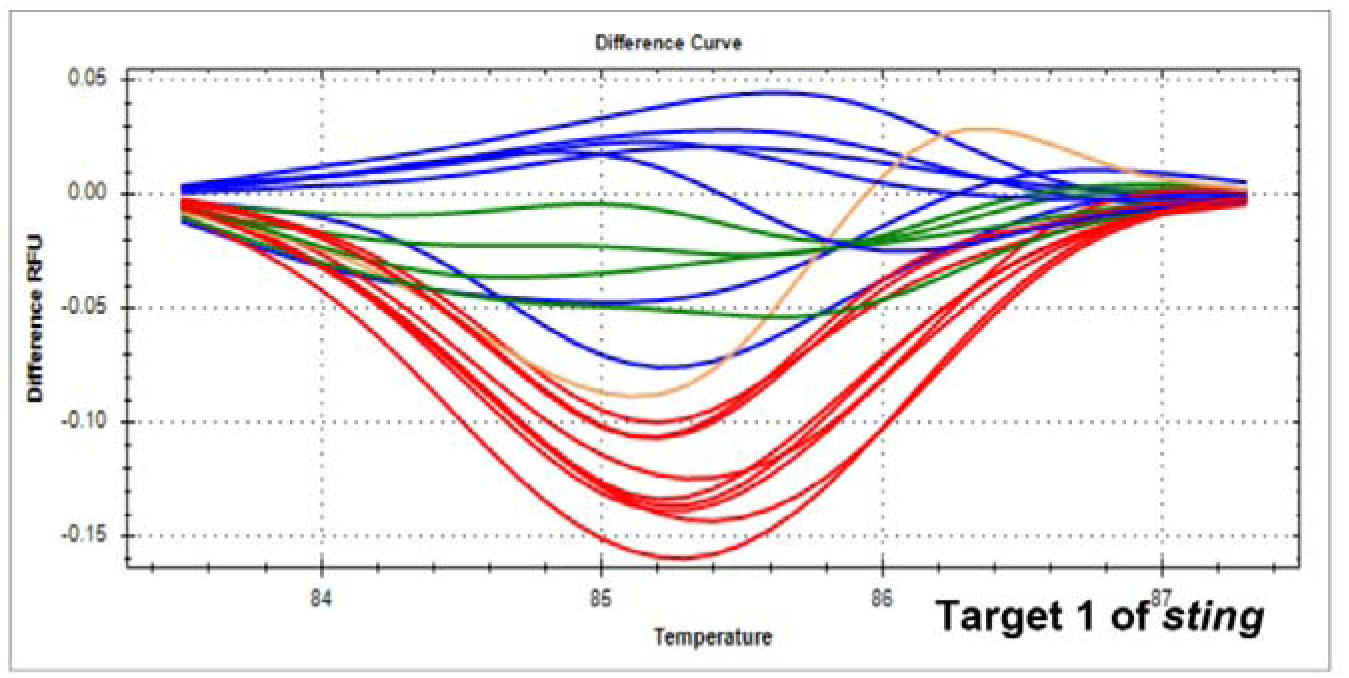
Editing efficiency of pooled gRNA-Cas9 editing of *sting*. High resolution melt analysis of *sting* gRNA1 target site, n=18, 77.77% efficiency, control clusters: Green & Blue.

**Supplementary Figure 4:**
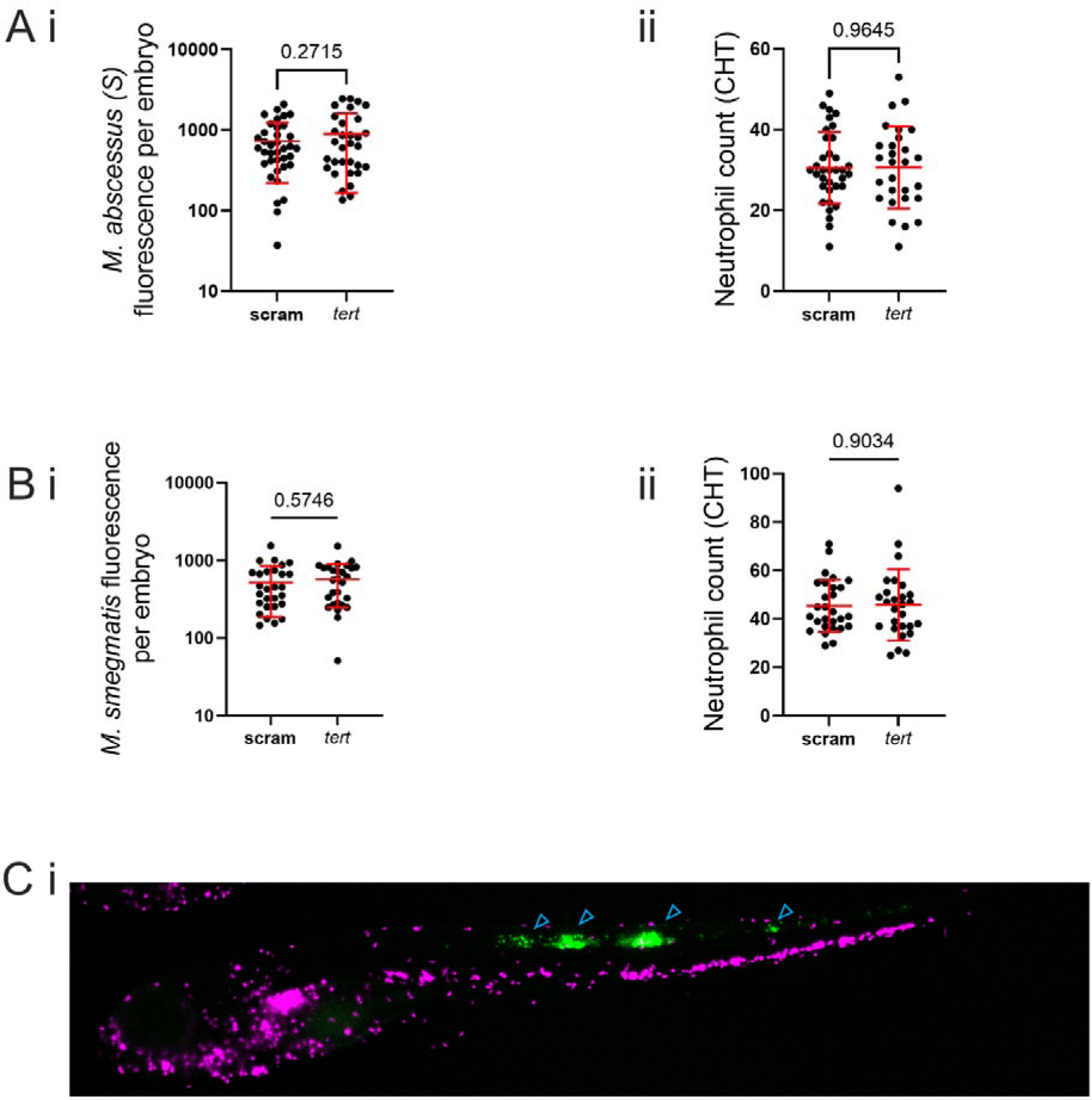
Tert-insensitive infection models do not induce demand-driven myelopoiesis (A) i. Quantification of smooth *M. abscessus* burden in 6 dpi *tert* crispant embryos. ii. Quantification of CHT-resident neutrophils in 6 dpi *Tg(lyzC:dsred) tert* crispant embryos infected with smooth *M. abscessus*. (B) i. Quantification of *M. smegmatis* burden in 6 dpi *tert* crispant embryos. ii. Quantification of CHT-resident neutrophils in 6 dpi *Tg(lyzC:dsred) tert* crispant embryos infected with *M. smegmatis*. (C) Representative image of neutrophils (magenta) and CHIKV-infected cells (green, indicated by blue arrowheads) in a 3 dpi *Tg(lyzC:dsred)* embryo.

