## Supplementary File for "Telomerase reverse transcriptase is required for resistance to mycobacterial infection"

| Use | Name | Sequence 5’-3’ | Reference/Note |
| --- | --- | --- | --- |
| Full length WT *tert* mRNA overexpression | FL tert | atgtctggacagtactcgacagatggcggatttaggccggttttggagattctgcgctccttatatccagtcgtgcagactttggaggagttcaccgacggactgcaattccctgacggccgaaagccggttctgctggaggaaacagacggcgcgcgctttaaaaagctcctcagtggacttattgtatgtgcgtacacgccgccgcagctgcgcgtccccgcccagctcagcaccctgccggaggtcttggcgttcactctgaaccacattaaacgtaagaaactgaggaacgtcctgggcttcggttatcaatgcagcgacgtgacgaccagttcggatcccttccgtttccatggcgacgtttcgcagacggctgcctccatcagcaccagcgaggtctggaagcgtatcaaccagcgtctgggcacggaggtaacgcggtacctgctgcaggactgtgccgttttcaccaccgtcccgccatcgtgtgttctgcaggtgtgcggagaacctgtttacgacttgctgatgccgcgctcatggtctggctttttcctcagtaactcagataatgaacgaatcagcggcgcgatgcggaaattccctgctgtccagaagacagtcgcaatttccaaaaagagaacaagagataacgaaaaatatatttcggtaaagcggcggagggtaaaggaaactgtgaataataataacggaaattacagatctctgtgttttgcaatttctaaaaagagagcgatagataatgaagaaaatatttcgttaaagcgacggaggatggaggaaactgaccaagtagcgaaaatacgtaatgaaaatcacgaatctcagagtttcgcaatttctaaaaagagagcgagagataatgaagaaaatatttcgttaaagcgacaaaggatggaggaaattgaccaagtagcgaaaatacgtaacgaaaatcatggatctcagagttggaaaccagcagatcagcgtcctcctcgaccctcgcaatgttcaatacgcgttctgagcatgctctacaatgggcggggcatgaagaacttcctgctcaacaggaagttgaaaggagtgggcggggccaggcgcatgcaaggggaggatcttgtccgcatgattttcctccaatcagaatccaacgacagcaaaccgaaaaaacttcccaaacgattcttcgcaatggtgccgctattcagtcggctgttgcggcagcacaggaagtgtccgtatcggctgttcctgcagaggaagtgtgcaggaaatccagacgtgaaggatatggagtctctgctgaagtcacactcgtctccatatagagtttatctgttcgtcagggagtgtctgcgccatattattccccacgagctctggggctgccaggaaaaccagctccacttcctgtctaatgtaaagaacttcctgcttctggggaagtttgagcgcctcacgctggtccagctgatgtggaggatgaaggttcaggcctgccattggctggggcccaagaaacgtcagtgtgcgagcgagcaccgctaccgtgagtggatgttgggtcagtgtatgggctggatgttgagtggttttgtggtcggtctggtcagagctcagttctacatcacggagagtatgggccacaaacacacactgcgcttctacaggggagatgtctggagcagactgcaggaccaggccttcagggctcatctgtgtaagggccagtggaggcccctgtctccatcccaggcgctgaaggtccccaatagtgcagtgacatcccgcatccgctttattcccaaaaccagcagcatgaggcccatcacacgcctcagcggcagcagagacacactgcagtattttcagagctgtgtgcgtgtgctgcagaatgtgttgagtgtgtgtgtgcgtgaggccccggggcccatgggctccaccgtctggggttggcaggacattcacagacgcctgcaagacttcagccctcagcagaagagctcgccacgaccgctctacttcgtcaaggtggatgtgagcggagcgtatgacagtctcccgcacctgaagctggtggaggtgctgaaggaagtgttgggtccgtttgcagagcagagcttcttcctgcgtcagtacagcagtgtgtggagcgacccgacccgcggcctgcgcaaacgcttctgcaccaaagctgagatgtcagagccgctcaacatgaaggggtttgttgtggatgaacaggtcagcgggcgcctgcatgacgctatattagtggagcggcactcgtctgaggtcagaggtggagacgtcttccagttcttccagaagatgctctgcagttacgtcatccattacgaccagcagatgttccggcaggtgtgtgggatcccgcagggctcttcagtgtcttctctgctgtgtaatctgtgttacggacacatggagaaagccctgctgaaggacatcgctaaaggagggtgtctgatgaggctgattgatgattttttgctcattactcctcatctgagtaaagccacagagttcctgaccactcttctgtctggagttccagattacggttgccagattaaccctcagaaggtggcggtgaacttccccgtgtgtgtgtcctgggtaaactcgggcgtctctgtgctgccgtccagctgcctgttcccctggtgcggcttgatgatacacacacacacgctggacgtctataaagactactcacggtatgacggcctatcactgcgctacagcctgactcttggctccgcccactctccatctacagtcatgaagaagctgctgtcggtgctcagcatcaaaagcacggacatcttcttagacctcaggctgaactctgtggaggccgtttacaggagtctgtataagctgattctgctgcaggcgctcaggtttcatgcgtgcgtgaggagtctgccgttgggtcagagtgtgaacagaaacccgtcgttcttcctgaagatgatctggagaatgactcgagtcaccaataaactcctcacacacattaacaaaggtctgcctgtgtgttctgtggacagtggtggtgttctgcagtctgaggcggttcagcttttattctgtttggccttcgagacgcttttcagacggtttcgctcggtttaccactgcctgatccctgcactgcacaaacggaagcgtgctcttcagcgtgagctctgcgggatcactctggctcgggtccgtcaagcttcctctcccagaatccccctggatttcagcatgcgggtgtaa |  |
| Full length ‘VAA’ *tert* mRNA overexpression | VAA tert | atgtctggacagtactcgacagatggcggatttaggccggttttggagattctgcgctccttatatccagtcgtgcagactttggaggagttcaccgacggactgcaattccctgacggccgaaagccggttctgctggaggaaacagacggcgcgcgctttaaaaagctcctcagtggacttattgtatgtgcgtacacgccgccgcagctgcgcgtccccgcccagctcagcaccctgccggaggtcttggcgttcactctgaaccacattaaacgtaagaaactgaggaacgtcctgggcttcggttatcaatgcagcgacgtgacgaccagttcggatcccttccgtttccatggcgacgtttcgcagacggctgcctccatcagcaccagcgaggtctggaagcgtatcaaccagcgtctgggcacggaggtaacgcggtacctgctgcaggactgtgccgttttcaccaccgtcccgccatcgtgtgttctgcaggtgtgcggagaacctgtttacgacttgctgatgccgcgctcatggtctggctttttcctcagtaactcagataatgaacgaatcagcggcgcgatgcggaaattccctgctgtccagaagacagtcgcaatttccaaaaagagaacaagagataacgaaaaatatatttcggtaaagcggcggagggtaaaggaaactgtgaataataataacggaaattacagatctctgtgttttgcaatttctaaaaagagagcgatagataatgaagaaaatatttcgttaaagcgacggaggatggaggaaactgaccaagtagcgaaaatacgtaatgaaaatcacgaatctcagagtttcgcaatttctaaaaagagagcgagagataatgaagaaaatatttcgttaaagcgacaaaggatggaggaaattgaccaagtagcgaaaatacgtaacgaaaatcatggatctcagagttggaaaccagcagatcagcgtcctcctcgaccctcgcaatgttcaatacgcgttctgagcatgctctacaatgggcggggcatgaagaacttcctgctcaacaggaagttgaaaggagtgggcggggccaggcgcatgcaaggggaggatcttgtccgcatgattttcctccaatcagaatccaacgacagcaaaccgaaaaaacttcccaaacgattcttcgcaatggtgccgctattcagtcggctgttgcggcagcacaggaagtgtccgtatcggctgttcctgcagaggaagtgtgcaggaaatccagacgtgaaggatatggagtctctgctgaagtcacactcgtctccatatagagtttatctgttcgtcagggagtgtctgcgccatattattccccacgagctctggggctgccaggaaaaccagctccacttcctgtctaatgtaaagaacttcctgcttctggggaagtttgagcgcctcacgctggtccagctgatgtggaggatgaaggttcaggcctgccattggctggggcccaagaaacgtcagtgtgcgagcgagcaccgctaccgtgagtggatgttgggtcagtgtatgggctggatgttgagtggttttgtggtcggtctggtcagagctcagttctacatcacggagagtatgggccacaaacacacactgcgcttctacaggggagatgtctggagcagactgcaggaccaggccttcagggctcatctgtgtaagggccagtggaggcccctgtctccatcccaggcgctgaaggtccccaatagtgcagtgacatcccgcatccgctttattcccaaaaccagcagcatgaggcccatcacacgcctcagcggcagcagagacacactgcagtattttcagagctgtgtgcgtgtgctgcagaatgtgttgagtgtgtgtgtgcgtgaggccccggggcccatgggctccaccgtctggggttggcaggacattcacagacgcctgcaagacttcagccctcagcagaagagctcgccacgaccgctctacttcgtcaaggtggatgtgagcggagcgtatgacagtctcccgcacctgaagctggtggaggtgctgaaggaagtgttgggtccgtttgcagagcagagcttcttcctgcgtcagtacagcagtgtgtggagcgacccgacccgcggcctgcgcaaacgcttctgcaccaaagctgagatgtcagagccgctcaacatgaaggggtttgttgtggatgaacaggtcagcgggcgcctgcatgacgctatattagtggagcggcactcgtctgaggtcagaggtggagacgtcttccagttcttccagaagatgctctgcagttacgtcatccattacgaccagcagatgttccggcaggtgtgtgggatcccgcagggctcttcagtgtcttctctgctgtgtaatctgtgttacggacacatggagaaagccctgctgaaggacatcgctaaaggagggtgtctgatgaggctgattgccgcctttttgctcattactcctcatctgagtaaagccacagagttcctgaccactcttctgtctggagttccagattacggttgccagattaaccctcagaaggtggcggtgaacttccccgtgtgtgtgtcctgggtaaactcgggcgtctctgtgctgccgtccagctgcctgttcccctggtgcggcttgatgatacacacacacacgctggacgtctataaagactactcacggtatgacggcctatcactgcgctacagcctgactcttggctccgcccactctccatctacagtcatgaagaagctgctgtcggtgctcagcatcaaaagcacggacatcttcttagacctcaggctgaactctgtggaggccgtttacaggagtctgtataagctgattctgctgcaggcgctcaggtttcatgcgtgcgtgaggagtctgccgttgggtcagagtgtgaacagaaacccgtcgttcttcctgaagatgatctggagaatgactcgagtcaccaataaactcctcacacacattaacaaaggtctgcctgtgtgttctgtggacagtggtggtgttctgcagtctgaggcggttcagcttttattctgtttggccttcgagacgcttttcagacggtttcgctcggtttaccactgcctgatccctgcactgcacaaacggaagcgtgctcttcagcgtgagctctgcgggatcactctggctcgggtccgtcaagcttcctctcccagaatccccctggatttcagcatgcgggtgtaa | This Study |
| Truncated C terminus domain *tert* mRNA overexpression | CTD tert | atgggctccgcccactctccatctacagtcatgaagaagctgctgtcggtgctcagcatcaaaagcacggacatcttcttagacctcaggctgaactctgtggaggccgtttacaggagtctgtataagctgattctgctgcaggcgctcaggtttcatgcgtgcgtgaggagtctgccgttgggtcagagtgtgaacagaaacccgtcgttcttcctgaagatgatctggagaatgactcgagtcaccaataaactcctcacacacattaacaaaggtctgcctgtgtgttctgtggacagtggtggtgttctgcagtctgaggcggttcagcttttattctgtttggccttcgagacgcttttcagacggtttcgctcggtttaccactgcctgatccctgcactgcacaaacggaagcgtgctcttcagcgtgagctctgcgggatcactctggctcgggtccgtcaagcttcctctcccagaatccccctggatttcagcatgcgggtgtaa | This Study |
| *egfp* mRNA overexpression as control | egfp | atgagtaaaggagaagaacttttcactggagttgtcccaattcttgttgaattagatggtgatgttaatgggcacaaattttctgtcagtggagagggtgaaggtgatgcaacatacggaaaacttacccttaaatttatttgcactactggaaaactacctgttccatggccaacacttgtcactactttctcttatggtgttcaatgcttttcaagatacccagatcatatgaaacggcatgactttttcaagagtgccatgcccgaaggttatgtacaggaaagaactatatttttcaaagatgacgggaactacaagacacgtgctgaagtcaagtttgaaggtgatacccttgttaatagaatcgagttaaaaggtattgattttaaagaagatggaaacattcttggacacaaattggaatacaactataactcacacaatgtatacatcatggcagacaaacaaaagaatggaatcaaagttaacttcaaaattagacacaacattgaagatggaagcgttcaactagcagaccattatcaacaaaatactccaattggcgatggccctgtccttttaccagacaaccattacctgtccacacaatctgccctttcgaaagatcccaacgaaaagagagaccacatggtccttcttgagtttgtaacagctgctgggattacacatggcatggatgaactatacaaataa |  |
